# Hypertrophic bone proliferation at enthesis induced by unilateral injection of botulinum toxin in masticatory muscles in adult rats is characterized by chondrocyte proliferation without inflammatory process at enthesis

**DOI:** 10.1101/2025.10.03.680271

**Authors:** Morgane Mermet, Quentin Massiquot, Nadine Gaborit, Stéphanie Lemiere, Jean-Daniel Kün-Darbois, Hélène Libouban

**Affiliations:** Univ Angers, Nantes Université, Oniris, Inserm, RMeS, REGOS, SFR ICAT, F-49000 Angers, France; Univ Angers, SFR ICAT F-49000 Angers, France; CHU Angers, Département de pathologie cellulaire et tissulaire, F-49933 Angers, France; CHU Angers, Service de Chirurgie Maxillo-faciale, F-49933 Angers, France

**Author notes:** **Correspondance to:** Pr Hélène Libouban, Inserm UMR_S 1229 RMeS – REGOS Team, University of Angers, Institut de Biologie en Santé, 4 rue Larrey, F-49933 Angers, France.

**Keywords:** Botulinum toxin, disuse osteoporosis, mandible, enthesis, masticatory muscle, inflammation

## Abstract

A single injection of botulinum toxin injected in masticatory muscles induces alveolar bone loss associated with muscle enthesis hypertrophic bone proliferation. The tissular and cellular mechanisms involved in this phenomenon are unknown. Because new bone formation due to inflammation at enthesis is observed in articular disorder like spondyloarthritis, we hypothesize that an inflammatory process could explain the development of hypertrophic bone proliferation in the BTX model. The aims of the study were to test this hypothesis and to determine whether IL-17A was involved.

Mature Sprague-Dawley male rats (n=36) were randomized into 4 groups. Three groups received BTX injections in right masseter and temporalis muscle. One BTX group was treated with an anti-IL17A antibody for 31 days (BTX31+antiIL17A). The two other groups were respectively sacrificed at 21 (BTX21) and 31 (BTX31) days post BTX injection. The remaining non-BTX group received anti-IL17A treatment for 31 days (control+anti-IL-17A). Alveolar bone loss and hypertrophic bone proliferation at enthesis were studied using microcomputed tomography. The presence of inflammatory process and the implication of IL17A was assessed by histology and immunohistochemistry on decalcified right side hemimandibles.

Quantitative measurement of the hypertrophic bone (bone volume and thickness) showed no significant differences between the 3 BTX groups. Histology revealed the presence of chondrocytes in large hypertrophic area but no inflammatory infiltrated cells. Immunochemistry confirmed the absence of inflammatory process but positive reaction for Ki67 which is in favor of actively growing and proliferative chondrocytes.

BTX-related muscle atrophy of masticatory muscle induced chondrocyte proliferation at enthesis without inflammation process.

## 1. Introduction

The use of botulinum toxin (BTX) to induce muscle atrophy is widely used to study the impact of low strain on bone. This model was originally developed to study disuse osteoporosis of hind limb and musculo-skeletal interactions [1]. In this model, BTX injection is performed in the *Mus. Quadriceps Femoris* of rat or mouse resulting in tibial and femoral bone loss at both cortical and trabecular level due to reduced mechanical strains [2-5]. The model was then reproduced in the mandible in which mandibular bone loss occurred one month after BTX injection [6]. In this model, injections of BTX in the two major masticatory muscles (temporal and masseter) induced a severe amyotrophy. Interestingly, an area of hypertrophic bone made of metaplastic woven bone at the accessory *Mus. Digastricus* enthesis was also described.

This phenomenon was systematically observed in the hemimandible where the injection of BTX was done (right hemimandible); the bone proliferation was absent in the left hemimandible used as control. Moreover, hyperproliferation of bone at enthesis is maintained after repeated unilateral injections of BTX in masticatory muscle [7].

The entheseal new bone formation can be compared to the fracture repair process in which an initial inflammatory phase precedes a mesenchymal tissue response with apposition of woven bone [8].

We hypothesized that the pathophysiology of the neoformation at the enthesis may be due to the increase of mechanical strains acting on the accessory muscle *Mus. Digastricus* to compensate the loss of activity of the two principal masticatory muscles (*Mus. Temporalis* and *Mus. Masseter*). This hypothesis is also in agreement with a previous study showing, in a model of spondyloarthritis (SpA), that enthesitis and new bone formation is driven by mechanical strain [9].

Moreover, we showed through analysis of computed tomography images, a bone apposition at the enthesis of *Mus. Digastricus* in patients treated with BTX, when this toxin was administered in the two muscles *Mus. Temporalis* and *Mus. Masseter* [10]. Thus, there is clear evidence that our animal model represents a model of metaplastic bone proliferation at enthesis due to mechanical stresses. It could though represent a good model to study the pathophysiological features of the tissue response consecutive to enthesitis.

Enthesitis is a main feature of spondyloarthritis. Excessive bone destruction and defective bone repair are typical features of rheumatoid arthritis (RA), while in ankylosing spondylitis (AS) there is excessive ectopic ossification and in psoriatic arthritis (PA) the two processes coexist. In all these diseases, increased interleukin-17 (IL-17) contributes to bone involvement [11]. New bone formation would represent a tissue response process that followed the optimal peak of inflammation at the enthesis.

The origin of the entheseal inflammation is still unclear and may involve several different processes. In SpA, entheseal inflammation could be explained by repeated mechanical stress or injury resulting in new bone formation [8]. One of the key mediators of inflammation at the enthesis is IL-23 [12] that activates Th17 cells to produce IL-17A and IL-22, responsible of inflammation, bone loss and ossification [13]. More specifically, IL-17A is suspected to increase and amplify inflammation at the enthesis and around [14]. In addition, the differentiation of human mesenchymal cells into osteoblasts is stimulated by IL17A [15] suggesting its implication in bone formation at the enthesis.

Several inhibitors of IL-17A were developed for the treatment of rheumatic and inflammatory diseases. Studies that involve established AS patients followed up over 2–4 years showed no evidence of the inhibitory effect of TNFi drugs on radiographic progression [16-18]. In contrast, a clinical 2 years study with secukinumab, an anti-IL-17A, suggests a low mean progression of spinal radiographic changes [19].

Human data concerning enthesitis are limited. Nevertheless, there are several animal models (mainly transgenic) of SpA. Although enthesitis is observed in these models, it is not associated to the formation of new bone [20]. However, the effect of an IL-17A vaccine on decreasing bonne destruction and inducing new bone formation in a transgenic rat model has been recently reported [21].

To date, all drugs used to treat enthesitis aim to stop inflammation and relieve symptoms. Knowledge on the enthesitis is limited especially because the concept that resolution of entheseal inflammation might also affect the new bone formation response is not well investigated [8].

Because of the absence of investigation of the mechanism explaining the development of hypertrophic bone proliferation at enthesis and the existence of similarity regarding new bone formation observed at enthesis in SpA, we hypothesized that an inflammatory process could be involved.

The aims of the study were thus to: (i) determine whether bone proliferation at enthesis induced by injection of BTX into the masticatory muscles was related to an inflammatory process at the enthesis of the digastric muscle in rats and (ii) to determine whether the process of bone proliferation at enthesis in the BTX model was linked to IL-17A by evaluating the effect of treatment with an anti IL17-A antibody in the BTX model.

## 2. Material and Methods

### 2.1. Animals and experimental procedure

Animal care and experimental protocols were approved by the French Ministry of Research and were done in accordance with the institutional guidelines of the French Ethical Committee (protocol agreement number APAFIS#15987-2018071012291155 v3) and under the supervision of authorized investigators.

Thirty-six male Sprague-Dawley mature rats aged 17/18 weeks and weighing 567 ± 34g were used for the study (Janvier-Labs, Le Genest-Saint-Isle, France). They were maintained under local vivarium conditions (24 °C and 12 h / 12 h light dark cycle) in the animal house facility of the University of Angers. Rats were given standard laboratory food (UAR, Villemoison-sur-Orge, France) and water *ad libitum*. The experimental procedure and analysis performed by samples type are summarized in Fig. 1.

**Figure 1:**
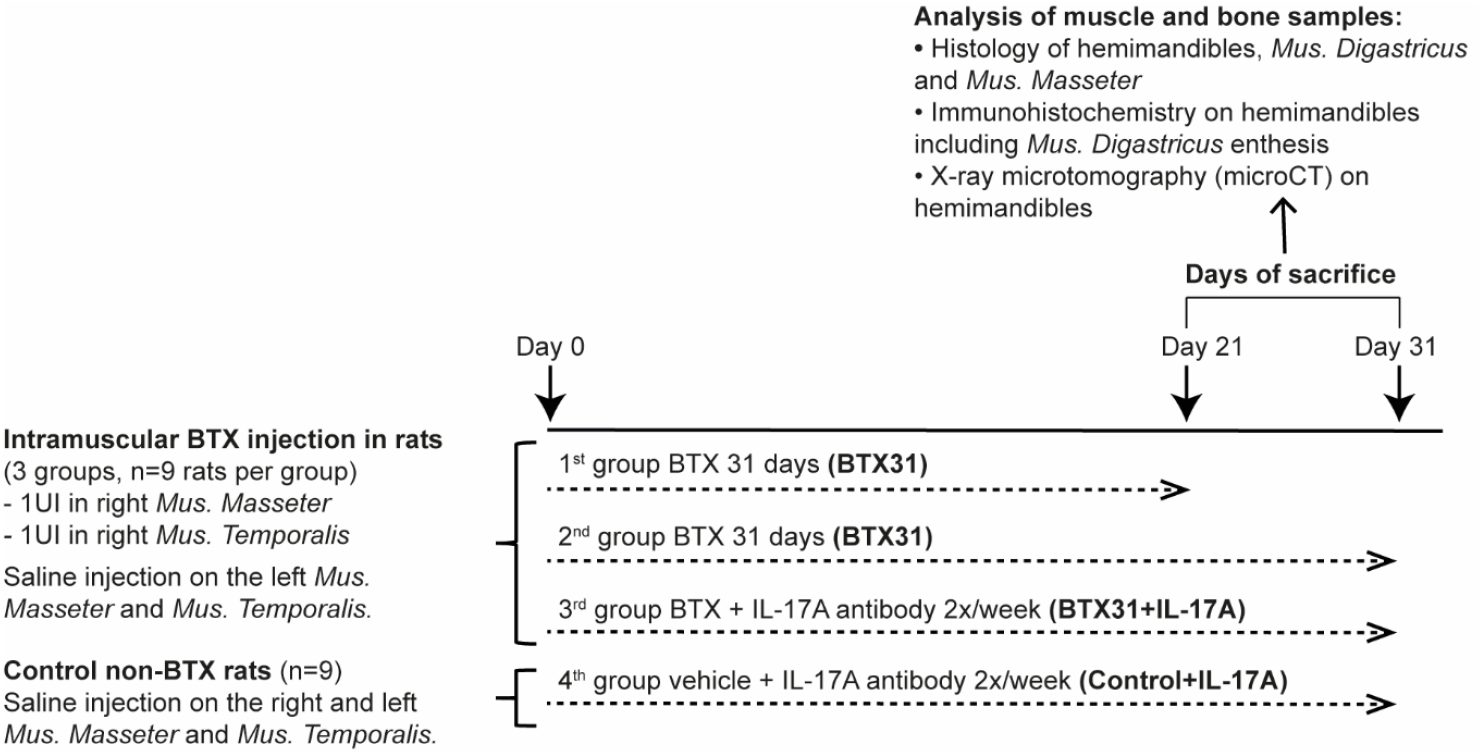
Experimental procedures and analysis performed by samples type.

At day 0, 27 rats were weighted, anesthetized with isoflurane and injected intramuscularly on the right side with 2U of type A BTX (Botox®, Allergan Inc., Irvine, CA, USA): 1U in the *Mus. Masseter* and 1U in the *Mus. Temporalis*. Three points of injection for each *Mus. Masseter* and two for each *Mus. Temporalis* were performed. The left non-injected side was considered as control.

At day 0, the remaining 9 rats were weighted, anesthetized with isoflurane and injected intramuscularly as previously described with saline vehicle (control non-BTX group). At day 0, a subset of BTX-injected animals (n=9) was treated with the mouse anti-IL-17A antibody (BZN035, from Novartis). We used the dose and the frequency of administration as recommended by Novartis: 15 mg / kg by intraperitoneal injection (*ip*), 2 times per week. Rats were weighed weekly and were sacrificed after injections by CO_2_ inhalation: at 21 days for a subset of BTX animals (n=9) and at 31 days for the remaining BTX animals and the two anti-IL-17A BTX and non-BTX groups.

The period of 31 days was necessary to generate significant bone proliferation at the *Mus. Digastricus* enthesis and mandibular bone loss as previously reported for this model [6]. The period of 21 days has been chosen to observe the starting period of ossification at the enthesis. Hemimandibles with the *Mus. Digastricus* enthesis were then dissected carefully and fixed in formalin until use. Mus. *Masseter* and *Mus. Digastricus* were also dissected and fixed in formalin until use.

### 2.2. X-ray microtomography (MicroCT)

MicroCT of right and left hemimandibles was performed using a Skyscan 1272 X-ray computerized microtomograph (Skyscan, Kontich, Belgium) equipped with an X-ray tube working at 70 kV / 142 µA. Bones were placed in plastic tubes filled with water to prevent desiccation. The tubes were fixed on a brass stub with plasticine and analyzed with an isotropic pixel size corresponding to 8.3 µm, the rotation step was fixed at 0.20°, the integration time fixed at 680 ms and exposure was done with a 0.5 mm aluminium filter.

For each hemimandible, a stack of 2D-sections was obtained and reconstructed using NRecon software (Bruker, version 1.7.1.0) and analyzed with CTan software (Skyscan, release 1.18.8.0).

#### 2.2.1. Enthesis hypertrophic bone proliferation measurements

2D and 3D measurements of the hypertrophic bone proliferation at the *Mus. Digastricus* enthesis were performed for each hemimandible using CTan software as previously described [7]. The following parameters were measured:

- Hypertrophic bone volume at enthesis (in mm^3^), by manually selecting the volume of interest from the first image of visible bone hypertrophy to the last image.
- Hypertrophic bone thickness at enthesis (in µm), which was defined as the longest height of bone proliferation at enthesis.
- Post enthesis cortical thickness (in µm), which was defined as the thickness of underlying cortical bone measured at the first distal section without visible bone proliferation.

On the left side, where no bone proliferation was expected, one measurement was performed on the corresponding section using anatomical reference points such as teeth roots, for equivalent post-enthesis cortical thickness.

#### 2.2.2. Alveolar bone measurements

Reorientation of the alveolar sections was performed using Dataviewer software (Bruker, version 1.5.6) to ensure the quality and reproducibility of 3D measurements. Alveolar bone measurements were done on 100 2D sections using CTan software. For each hemimandible, the section going through the middle of the 3 crowns was firstly selected and was defined as the middle section (Supplemental Fig. 1A). The anterior and posterior limits were determined 50 sections respectively below and above the middle section. Regions of interest for measurement were then manually selected to create a 3D volume of interest (VOI) (Supplemental Fig. 1B). Teeth roots, alveolar canal and cortical bone were not included in the VOI which comprised only alveolar bone. 3D alveolar bone volume (BV/TV_3D_) was obtained using CTan. According to guidelines for assessment of bone microarchitecture in rodents using MicroCT, BV/TV is the fraction of the VOI occupied by alveolar bone, expressed in % [22].

### 2.3. Histology

#### 2.3.1. Hemimandibles

Data for the hemimandibles were first monitored using microCT. Osteotomy was then performed to isolate the molar area before decalcification of the samples with 10% ethylenediaminetetraacetic acid (EDTA, pH 7.2), during 3 to 4 weeks. The specimens that include the metaplastic bone proliferation zone, the *Mus. Digastricus* enthesis and the extremity part of *Digastricus* muscle were then dehydrated, defatted and embedded in paraffin using an automate apparatus (Tissue-Tek VIP 6 AI). Serial frontal sections of 5 µm thickness were performed for standard histological staining and for immunochemistry.

For standard histology, sections were stained with HPS (Hematoxylin Phloxine Saffron) as a standard tissue stain for the microscopic observation of bone metaplastic at the enthesis and the presence of inflammatory cells.

#### 2.3.2. Mus. Masseter and Mus. Digastricus muscle

After 24 hours of fixation, muscles samples were transaxially cut in the middle of the muscle to obtain cross sections of muscles fibers. The cut samples were then dehydrated, defatted and embedded in paraffin (with the flat edge of the cut samples embedded down) using an automate apparatus (Tissue-Tek VIP 6 AI). Serial frontal sections of 5 µm thickness were performed for standard histological staining and for immunochemistry.

Sections were stained with HPS as a standard tissue stain for the microscopic observation of atrophic masseter muscle and for structural aspects of digastric muscle.

### 2.4. Immunohistochemistry (IHC)

The following proteins were identified using specific anti-rat primary antibodies: CD68 to analyze the macrophage population; CD177 to analyze the polynuclear neutrophil population; CD4 for T cells CD4+; CD8 for T cells CD8+; IL-17A as the main cytokine produced by Th17 and Ki67 as a marker of proliferating cells. All the antibodies were polyclonal and from rabbit host.

Hemimandible decalcified sections were deparaffinized and antigen retrieval was performed by Heat Induced Epitope Retrieval (HIER) in a citrate buffer (pH 6) using the Bio SB TintoRetriever Pressure Cooker (cat#BSB-7005, Santa Barbara, USA). The protocol of 20 minutes at 100°C, at atmospheric pressure, was applied. Endogenous peroxidases were then inactivated by incubating sections in H_2_O_2_ 3% for 10 minutes. The non-specific binding sites were blocked by incubating the sections with casein solution (blocking solution from Candor, Wangen, Germany) for 1 hour. The sections were then incubated with each primary antibody for 1 hour at room temperature. The reference and the dilution for each antibody are mentioned in table 1. After primary antibody incubation, the sections were rinsed 3 times with Tris Buffer Saline (TBS) before incubation for 30 minutes at room temperature with a horseradish peroxidase-conjugated secondary antibody for rabbit antibodies (Polink-1 HRP detection system from GBI Labs, Rockville, USA). After washing with TBS, the immunological reaction was revealed by incubation of sections with DAB (3,3’-diaminobenzidine) chromogen for 5 minutes.

**Table 1.**
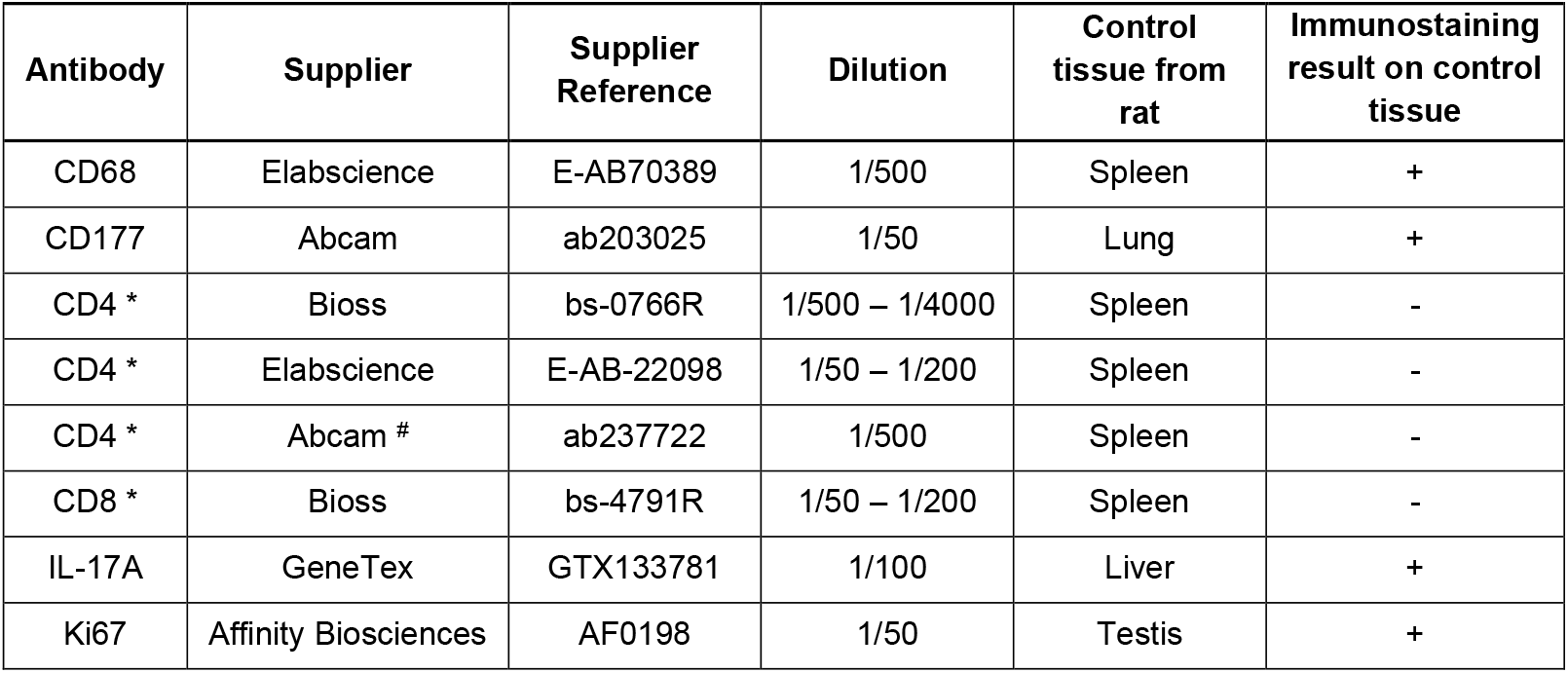
Antibody characteristics and validation for immunoreaction. * Antibody that we could not validate for immunostaining ; # Monoclonal antibody

Sections were then counterstained with Mayer’s hemalum solution for 20 seconds. Brown staining in cells was considered positive.

Sections from left and right side for each group of rats were carefully examined using optical microscopy at magnification 100X and 200X to analyze the immune reaction. A previous validation of all antibodies mentioned in Table 1 was performed from control tissues from rats before their use on the hemimandible sections. Those positive control tissues were determined according to the literature and to information mentioned in the sheet from the supplier of each antibody. The specific control tissue used for each antibody is presented in table 1. In case of negative or weak reaction, on those control tissue, several commercial antibodies from different suppliers were tested for a same antigen (like CD4. Table 1).

### 2.5. Statistical analysis

Statistical analysis was performed with GraphPad Prism 8.0.1 (GraphPaD software, La Jolla,CA, USA). All data were expressed as mean ± standard deviation (SD). One-way analysis of variance (ANOVA) with Tukey’s multicomparison test was used for enthesis hypertrophic bone proliferation measurements performed only on the right side. For the other measurements, two-way analysis of variance (ANOVA) was used: (i) to determine significant differences on weight between groups and between times; (ii) to determine significant differences on bone measurements between groups and between left and right sides with respectively Tukey’s or Sidak’s multiple comparison test. Differences at *p* < 0.05 were considered significant.

## 3. Results

### 3.1. Body weight and anatomic examination of maxillo-facial region

No significant difference in body weight was evidenced between groups at any time studied. However, significant differences in body weight were evidenced at time of sacrifice (21days for the BTX group or 31 days for the 3 other groups) *versus* day 0 (Fig. 2).

**Figure 2:**
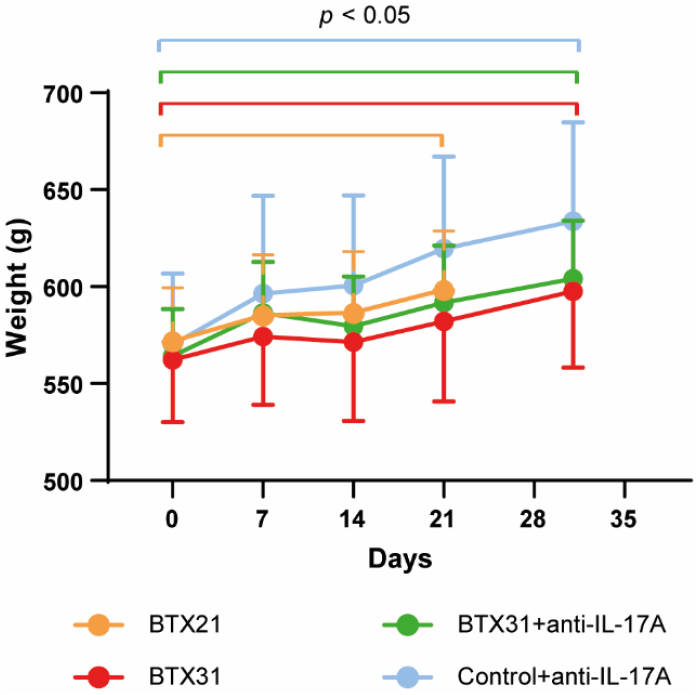
**Evolution of the body weight of rats** showing significant differences at 21 days for the BTX21 groups or 31 days for the 3 other groups (BTX31, BTX31+anti-IL-17A and Control+anti-IL-17A) versus day 0.

Visual examination during dissection revealed asymmetry of the incisors for 5 among the 9 rats in both BTX31 groups treated or not with IL-17A. No asymmetry was observed in the BTX21 group and in the control non-BTX group treated with IL-17A.

All rats of the three BTX groups revealed amyotrophy of the temporal and masseter muscles on the right side at 21 or 31 days post injection of BTX.

### 3.2. MicroCT analysis

#### 3.2.1 MicroCT analysis of enthesis hypertrophic bone proliferation measurements

As expected, hypertrophic bone proliferation at enthesis could only be observed on the right side of BTX injected animals. For the three BTX groups where bone proliferation at enthesis was expected, the number of rats presenting a hypertrophic bone at enthesis was, respectively: (i) 9 rats out of 9 for the BTX21 group; (ii) 9 rats out of 9 for the BTX31 group; (iii) 8 rats out of 9 for the BTX31+anti-IL-17A group.

No significant differences were observed between the three BTX groups concerning the hypertrophic bone volume and thickness at enthesis (Fig. 3A, B).

**Figure 3:**
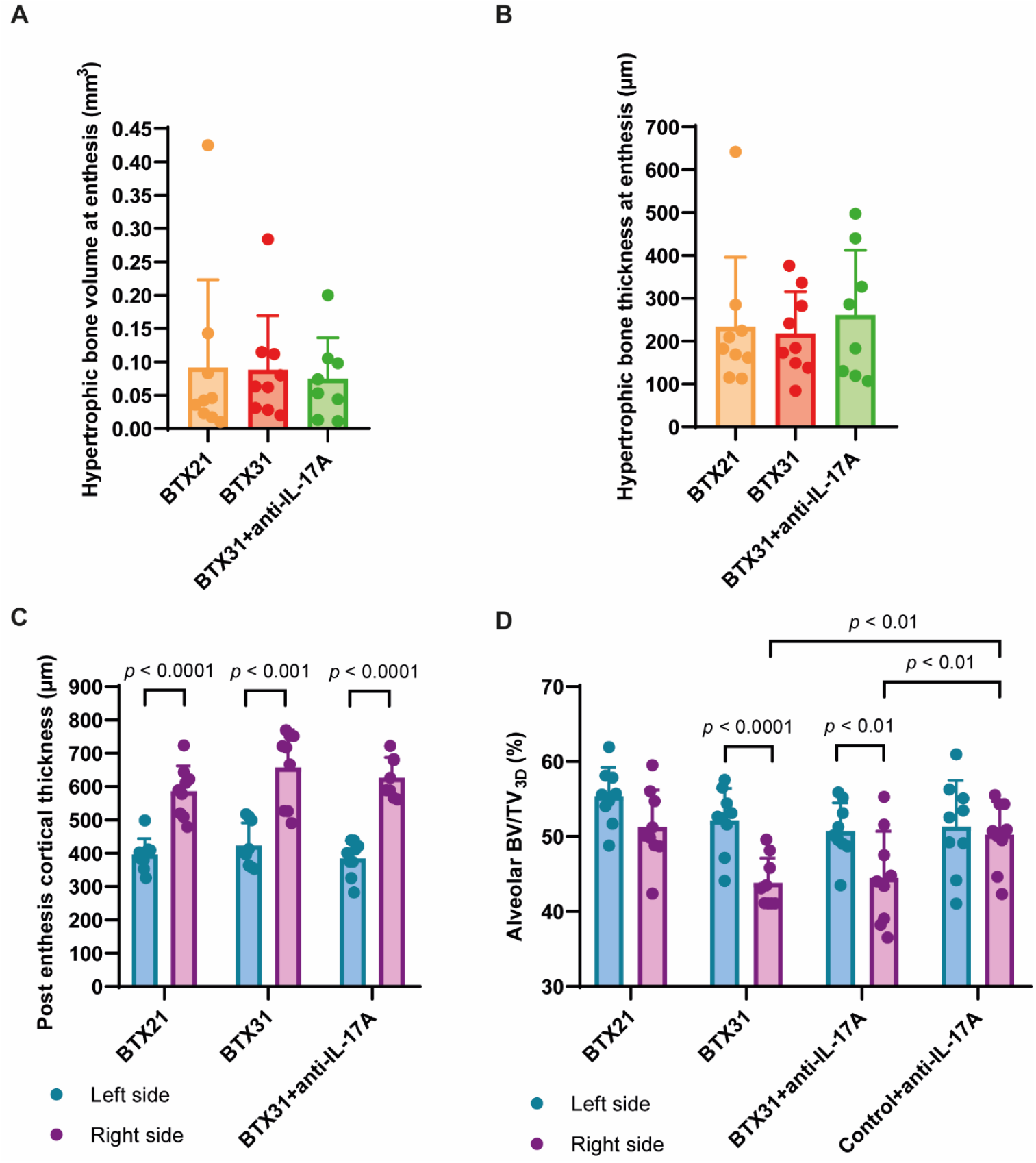
**Bone morphometric parameters** regarding hypertrophic bone (A, B), cortical bone post enthesis (C) and alveolar bone (D).

Post-enthesis cortical thickness was significantly higher on the right injected side compared to the left non-injected side in all the 3 BTX groups (Fig. 3C), supporting the animal model and the correct response to BTX injection. No difference was observed when considering the post-cortical thickness between the right side of the 3 BTX groups.

#### 3.2.2. MicroCT analysis of alveolar bone measurements

Alveolar BV/TV_3D_ was significantly reduced on the right injected side compared to the left non-injected side in BTX31 and BTX31+anti-IL-17A groups (Fig. 3D). A trend of reduction of alveolar BT/TV_3D_ was observed in the BTX21 group with no significance. On the right side, BV/TV3D was significantly higher in the control non-BTX group treated with anti-IL-17A compared to BTX31 and BTX31+anti-IL-17A groups. No difference was observed when considering the alveolar BV/TV_3D_ between the left side of the 4 groups.

### 3.3. Histological observation of hemimandibles and muscles

#### 3.3.1. Hemimandibles

General histological view at low magnification (50X) of the hypertrophic bone region on the right side is presented in Fig. 4A compared to the control left side. Histological view at low magnification (50X) from the control left side showed the absence of hypertrophic bone in close proximity of hemimandibles cortical bone (Fig. 4A – double black arrow), a tendinous insertion (Fig. 4A - T) into hemimandibles cortical bone and the attachment of muscle fibers to the cortical hemimandible from the tendon.

**Figure 4:**
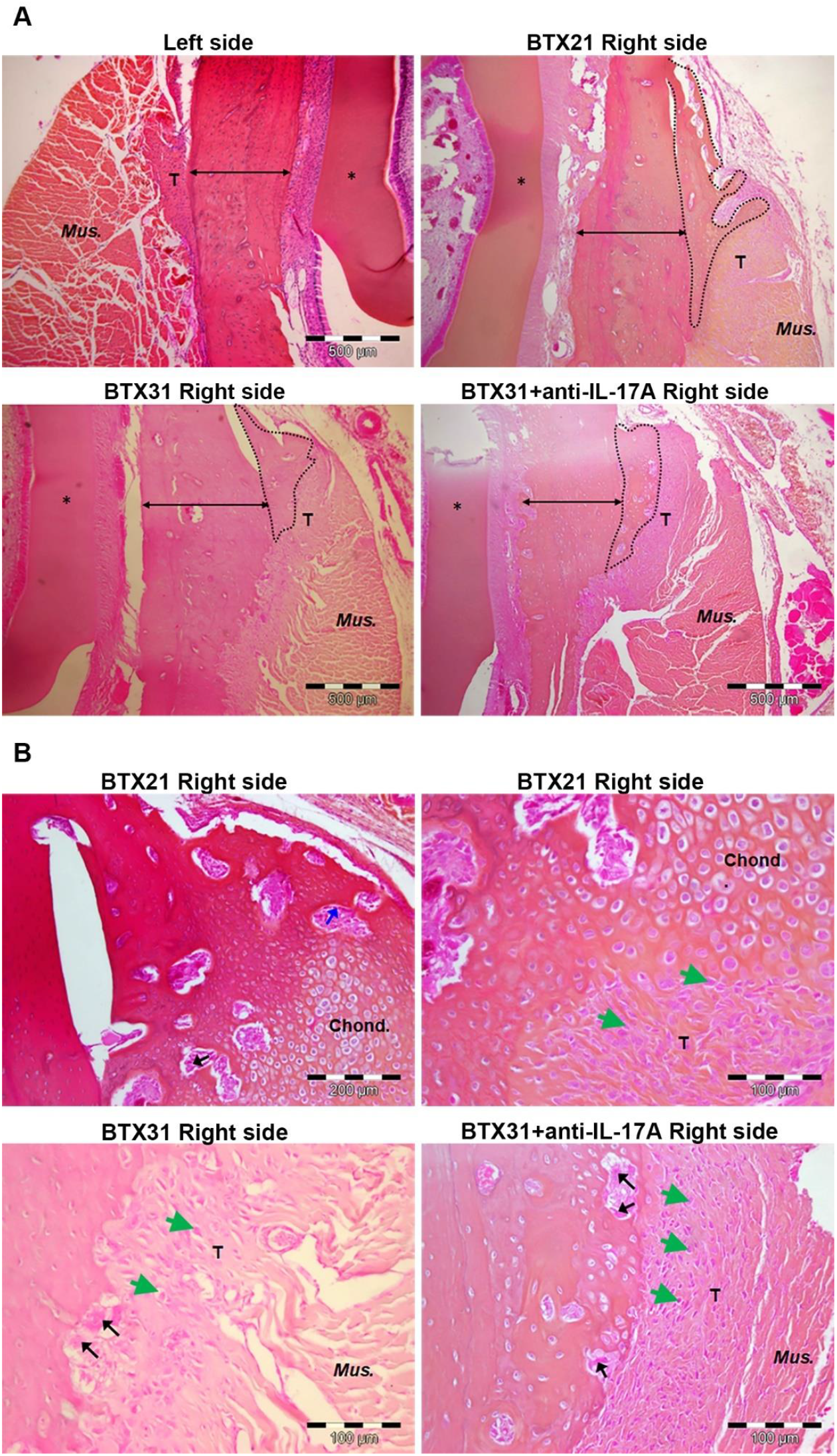
**Histological observation of hypertrophic bone region at enthesis** with low magnification (A) and high magnification (B). A: Hypertrophic bone region outlined by black dotted line / Cortical bone delimited by black arrow / Mus.: *Mus. Digastricus* / T: Tendon / *: dentine of rat incisor. B: Black arrows designating active osteoclasts / Blue arrow designating osteoblast alignment / Green arrowheads designating cells with a chondrocyte appearance / Mus.: *Mus. Digastricus* / T: Tendon.

In contrast, on the right side of the 3 BTX groups, hypertrophic bone was evidenced (Fig. 4A – bounded by a black dotted line). Separation between normal cortical bone (Fig. 4A – double black arrow) and hypertrophic bone was clearly identified. Tendinous regions appeared larger with a metaplastic appearance.

Small or large hypertrophic bone areas were observed in all the 3 BTX groups. We could not observe a trend of smaller hypertrophic areas in the BTX 21 group compared to the 31BTX groups with or without anti-IL-17A treatment.

Histological observations at higher magnification (100X - 200X) for the 3 BTX groups at the right side are presented in Fig. 4B. An active remodeling of the hypertrophic bone in all the BTX groups was observed. Osteoblasts alignments were evidenced as well as osteoclasts (Fig. 4B - blue and black arrows, respectively). In BTX31 and BTX31+anti-IL-17A, numerous eroded zones with large osteoclasts were observed at the periphery of hypertrophic bone as well at the periphery of cortical bone under the hypertrophic bone area.

We observed mostly in the BTX21 group and to a lesser extent in the BTX31 group, the presence of chondrocytes for the samples that presented large hypertrophic bone area. They appeared mostly packed like isogenous groups in cartilage (Fig 4B. 100X and 200X magnification). To confirm the presence of cartilage in specific samples with large hypertrophic bone area in the BTX21 group, a blue alcian staining was performed. It is commonly used to stain cartilage tissue as it detects cell chondrogenesis and sulfated proteoglycans. Cartilage appeared in blue at the interface with proliferative zone area and around active remodeling region (Supplemental Fig. 2).

A fibro-cartilaginous transition region was evidenced at the interface with the chondrocytes. This region presented some cells that did not have a fibroblastic aspect but rather an aspect close to that of chondrocyte (Fig. 4B - green arrowheads). Similar cells with a chondrocyte aspect in the metaplastic tendinous regions in both BTX31 groups were observed.

In the Control group+IL-17A, no histological modification was observed at the *Mus. Digastricus* enthesis either on the left or on the right side of the mandibles (data not shown).

#### 3.3.2. Mus. Masseter

The histological aspect of *Mus.Masseter* fibers at left and right side of the four groups is presented in Fig. 5A.

**Figure 5:**
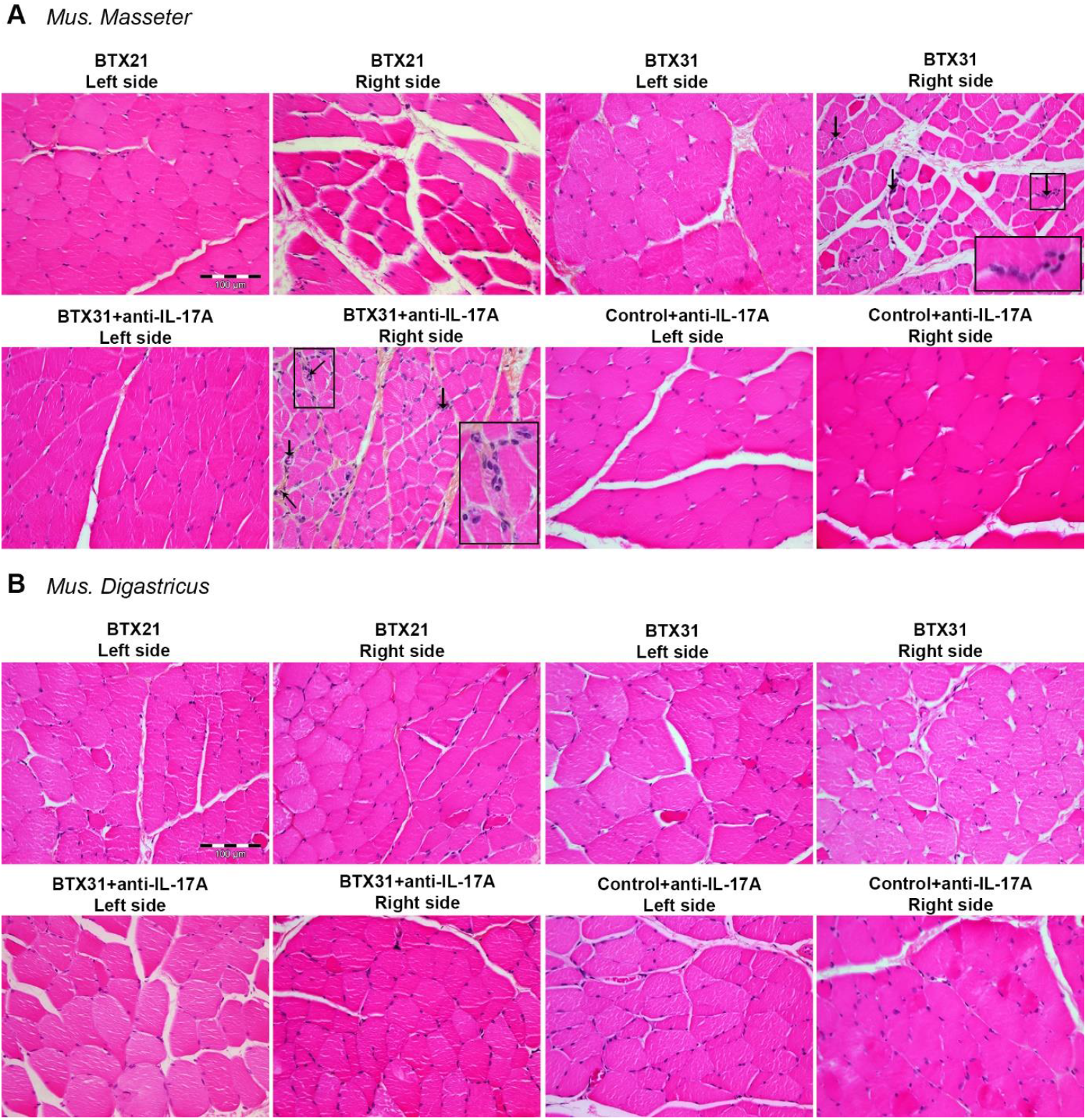
Histology of *Mus. Masseter* (A) and *Mus. Digastricus* (A). Black arrow represented infiltrated cells and were observed in both BTX31 groups.

Histological atrophy of the *Mus. Masseter* was evidenced on HPS sections 21 days after BTX injections. Compared to the left side, muscle fibers were less closely packed together and the conjunctive tissue (endomysium) that surrounded each muscle fiber was visible. 31 days after BTX injections, the histological alteration of the muscle fibers was amplified compared to the histological aspect at 21 days. Several fibers presented a cross-section surface smaller than the left side. The histological aspect of the *Mus. Masseter* in the BTX31+anti-IL-17A group was similar to that of the BTX31 group. Some infiltrated cells were also observed in the two BTX31 groups (Fig. 5A - black arrow).

In the control non-BTX group, no histological difference was observed between the left and the right *Mus. Masseter*.

No histological difference was observed between the left side non-injected muscle in the 4 groups.

#### 3.3.3 Mus. Digastricus

Histological aspect of *Mus. Digastricus* fibers at left and right side from the four groups is presented in Fig. 5B.

For each group, similar histological aspect was observed between the left and the right *Mus. Digastricus*. The muscle fibers were closely packed together with, sometimes, a thin visible endomysium. No sign of hypertrophy, nor infiltrated cells were evidenced

### 3.4 Immunohistochemistry

CD68, CD177, IL-17A and Ki67 antibodies were validated due to positive reaction on their specific control positive tissue (see “Immunostaining result on control tissue” column in table 1). For CD4 and CD8 immunostaining, antibodies could not be validated on spleen despite 3 antibodies tested from 3 different suppliers especially for CD4 and different dilution for both CD4 and CD8 antibodies (see dilution column in table 1).

For each immunoreaction on right hemimandibles, we analyzed the presence or absence of positive cells around the hypertrophic bone region including tendon with or without metaplastic aspect and *Mus. Digastricus* anchored to the tendon.

IHC reaction revealed an absence of immunostaining at the enthesis of *Mus. Digastricus* for CD68, CD177 and IL17A, regardless of the BTX groups.

In contrast, positive immunoreactions were observed in the 3 BTX groups for Ki67 antibody (Fig 6). Positive cells were localized in the close proximity of hypertrophic bone region. In BTX31 + IL-17A group, the positive staining was less extended with a less intense brownish staining in positive cells compared to positive sections in BTX21 and BTX31 groups. In BTX21 group, chondrocytes were also positive within the cartilaginous zone (outlined by black dotted line – Fig. 6). Some cells with a chondrocyte aspect were also positive (green arrowheads – Fig. 6).

**Figure 6:**
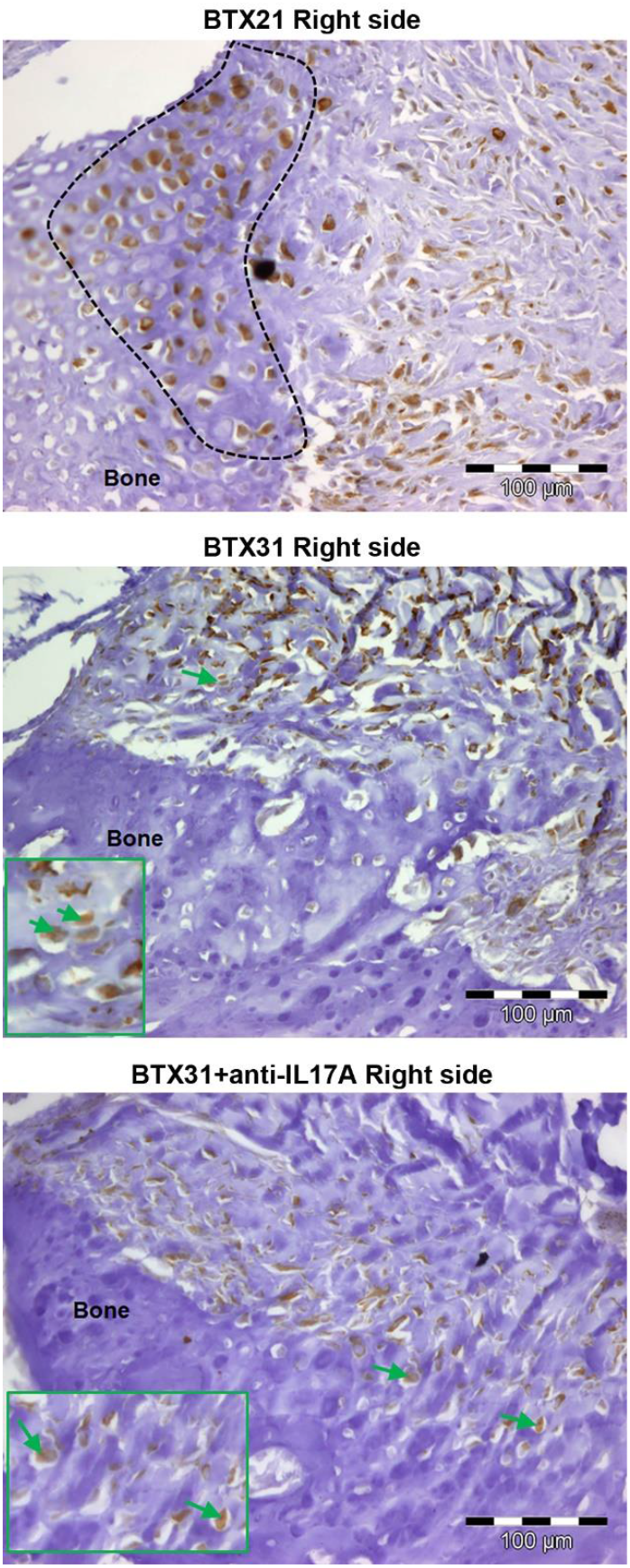
Immunohistochemistry for Ki67 at enthesis. Positive cells appeared in a brownish color. Black dotted line delimited chondrocyte positive cell for Ki67; green arrowheads designed positive cells with a chondrocyte aspect; “bone” referred to hypertrophic bone.

## 4. Discussion

One of the two objectives of the study was to determine if treatment with an anti IL17-A antibody in the BTX model was able to inhibit or reduce the hypertrophic bone observed at enthesis of *Mus. Digastricus*. MicroCT analysis allowed to answer this question.

We had planned to answer the question by determining the number of rats per group that developed or not hypertrophic bone at enthesis without volume consideration of enthesis. However, the absence of differences in the number of rats that developed or not hypertrophic bone at enthesis showed that this phenomenon occurred rapidly after the induced atrophy of masticatory muscle and earlier than the period of 21 days. Moreover, this result showed also that treatment with an anti IL17-A antibody has no effect on the development of hypertrophic bone at enthesis of *Mus. Digastricus*. Within the same BTX group (BTX21, BTX31 or BTX31+anti-IL-17A), we observed on microCT images a large heterogeneity in the size of the hypertrophic bone. Quantitative methods that we have proposed and applied in a recent study on repeated injection of BTX in masticatory muscle help to better characterize the hypertrophic bone at enthesis and confirm heterogeneity in term of the size of the hypertrophic bone within a same group [7].

Data on hypertrophic bone volume and thickness in the BTX31+anti-IL-17A group confirmed that treatment with an anti IL17-A antibody has no effect on the development of hypertrophic bone at enthesis of *Mus. Digastricus*.

We also hypothesized that bone volume at enthesis could be significantly lower at 21 days compared to 31 days after the BTX inoculation but again, large or small hypertrophic bone could be observed at 21 days as well as at 31 without any significant difference. The starting period of ossification at enthesis that we hypothesized at 21 days was not found reproducible between rats and should be earlier. Moreover, it is probably difficult to determine this period by microCT or histology at an earlier time (especially by histological methods). MicroCT would be easier, but it is a method that cannot give information on the cellular processes conducting to hypertrophic bone.

We also reported a thickening of the enthesis adjacent cortical bone in all BTX groups. This observation was recently confirmed after unilateral repeated injection of BTX in masticatory muscles [7]. However, the thickening of enthesis at adjacent bone cannot be explained and needs to be more investigated. We can hypothesize that a localized increase in muscle activity may lead to modification at *Mus. Digastricus* enthesis and also at the same time to changes in adjacent cortical bone. However, the histological examination of *Digastricus* muscle did not show signs of hypertrophy. Treatment with an anti IL17-A antibody has also no influence on the adjacent cortical bone.

We measured alveolar bone since it is known to decrease following BTX injection [6,7,23]. We confirmed an alveolar bone loss at the injected side after 31 days with a trend to a decrease at 21 days. The two specific times chosen for this study, 21 and 31 days, were therefore appropriate to characterize the bone loss consecutive to *Mus. Masseter* atrophy. Muscle loading plays an essential role in maintaining bone volume and density [24]. Mechanical forces exerted by the masticatory muscles stimulate the mandibular bone and alveolar process to maintain the underlying bone and teeth state. Alveolar bone mineral density is thus considered as correlated with masticatory function. Treatment with an anti IL17-A antibody has no influence on the bone alveolar loss. Moreover, it did not modify alveolar bone volume in the control non-BTX group.

Recent findings showed that IL-17 cytokines through IL17-RA signaling can stimulate osteoclasts and contribute to bone loss in osteoporosis characterized by an inflammatory etiology like osteoporosis due to estrogen deficiency [25,26].

Our results are in favor of a non-implication of IL-17A in alveolar bone loss in the BTX model which is in accordance with the literature in which no implication of IL-17A was reported in disuse osteoporosis.

The second objective of the study was to determine whether bone proliferation at enthesis induced by injection of BTX into the masticatory muscles was related to an inflammatory process at the enthesis of *Mus. Digastricus*. Histological examination of hemimandibles followed by immunohistochemistry allow to answer this question but also to better characterize the cellular and tissular modifications at enthesis of *Mus. Digastricus*.

Bone proliferation occurring in BTX rats has not been studied by other authors although the same phenomenon seems to occur in human after BTX injections [10]. Enthesis morphology seems to depend on the local mechanical environment, *i.e*., muscle activity, to which it is subjected.

No histological study was conducted on the hypertrophic bone region induced by BTX at enthesis. Histological organization of enthesis, its cellular component and its mechanical properties depends on its anatomical location [27]. Enthesis of *Mus. Digastricus* in the non-injected control side is fibrous with the tendon inserted between the muscle fibers and the hemimandibles cortical bone. In contrast, enthesis *of Mus. Masseter* is fibrocartilaginous. BTX injection induced severe histological modifications at enthesis including hypertrophic bone and an extend of the fibrous insertion with a metaplastic appearance.

Treatment with an anti-IL17A antibody has no influence on the cellular and tissular events at enthesis of *Mus. Digastricus*. Moreover, it has no effect on bone erosion that was abundant 31 days after BTX injection.

Bone erosion is one of the characteristics observed in spondyloarthritis and it is reduced by IL-17A inhibitors [28]. The fact that bone erosion is not reduced in the BTX31 group treated with IL-17A antibody means that IL-17A is not implicated in the activation of osteoclasts at the periphery of hypertrophic bone and at the periphery of the adjacent cortical bone.

An interesting finding of this study is the presence of numerous chondrocytes and an osteo-cartilaginous zone in the close proximity of hypertrophic bone in the BTX21 groups with large hypertrophic bone at enthesis. For fibrocartilaginous type enthesis, area of fibrocartilage decreases in unloading condition [29]. Moreover, in the BTX model, a decrease in fibrous chondrocytes at enthesis of *Mus. Masseter* was observed 6 weeks after BTX injection [30]. The apparition of fibrocartilaginous tissue around hypertrophic bone is in favor of the hypothesis of the increase of mechanical strain at enthesis of *Mus. Digastricus* to compensate the loss of activity in *Mus. Masseter* and *Mus. Temporalis*. Positive Ki67 immunostaining in chondrocytes observed in BTX21 groups and in the cells with a chondrocyte aspect in both BTX31 groups reinforce this mechanical hypothesis. It has thus recently been showed that Ki67 is a good marker of actively growing and proliferating chondrocytes [31]. Moreover, a decrease in chondrocyte proliferation is observed in the absence of mechanical strain [29].

Although histological examination of the enthesis showed numerous cells in the fibrous tendon, no infiltrated cells evoking inflammation were observed. Moreover, immunohistochemistry revealed an absence of macrophages and neutrophils by respectively negative staining for CD68 and CD177. Due to poor activity of commercial antibodies targeting CD4 and CD8, we could not identify without ambiguity the presence of T lymphocytes. Likewise, IL-17A was not implicated. Based on these results, it seems that there is no inflammation process implicated in the hypertrophic bone at enthesis of *Mus. Digastricus*. We could think that the times (*i.e*., 21 and 31 days) chosen for this study were not optimum to detect an inflammatory process prior to hypertrophic ossification. On the other hand, the lack of effect of IL-17A treatment is not in favor with an inflammation process to explain hypertrophic ossification.

In conclusion, the study showed that hypertrophic bone that developed at enthesis of *Mus. Digastricus* is not due to inflammation process through IL-17A pathway. Increase of mechanical strain in *Mus. Digastricus*, because of the absence of mechanical strain in principal masticatory muscles, seems to be the main mechanism of induced hypertrophic bone at enthesis. The study allowed to characterize the tissular and cellular modifications at enthesis *of Mus. Digastricus*: the fibrous enthesis evolves to a fibrocartilaginous enthesis around hypertrophic bone. Some questions remain about the cellular and molecular process involved in ossification as well as the process leading to metaplastic tendon which seem to be more impacted in its structure and morphology than *Mus. Digastricus* itself.

## Acknowledgements

The authors thank Novartis for the funding of the full study and for providing the mouse anti IL17-Antibody (BZN035). The author thanks Dr. Christelle Darrieutort-Laffite for her help in reviewing the manuscript. The authors thank the HiMolA Platform, University of Angers for assistance with bone evaluation. The authors are grateful to the SCAHU platform, SFR ICAT, University of Angers for animal housing and care.

## CRediT authorship contribution statement

**Morgane Mermet:** Investigation, Methodology, Formal analysis, Data curation; **Quentin Massiquot:** Investigation, Methodology, Formal analysis, Data curation; **Nadine Gaborit:** Investigation, Methodology, Formal analysis, Data curation; **Stéphanie Lemière:** Investigation, Methodology, Formal analysis, Data curation; **Jean Daniel Kün-Darbois:** Investigation, Methodology, Editing; Resources, **Hélène** Review & **Libouban:** Conceptualization, Formal analysis, Writing - Review & Editing, Supervision, Funding acquisition, Data curation

## Data availability statement

The data that support the findings of this study are available from the corresponding author upon reasonable request.

