## Supplemental Figures 1&2 for "Hypertrophic bone proliferation at enthesis induced by unilateral injection of botulinum toxin in masticatory muscles in adult rats is characterized by chondrocyte proliferation without inflammatory process at enthesis"

**Supplemental Fig. 1.**

MicroCT analysis for 3D alveolar measurements of BV/TV. The anterior and posterior limits were determined according to the red line that defined the middle section going through the 3 crowns (A). Region of interest used for alveolar 3D measurement appeared in red (B).

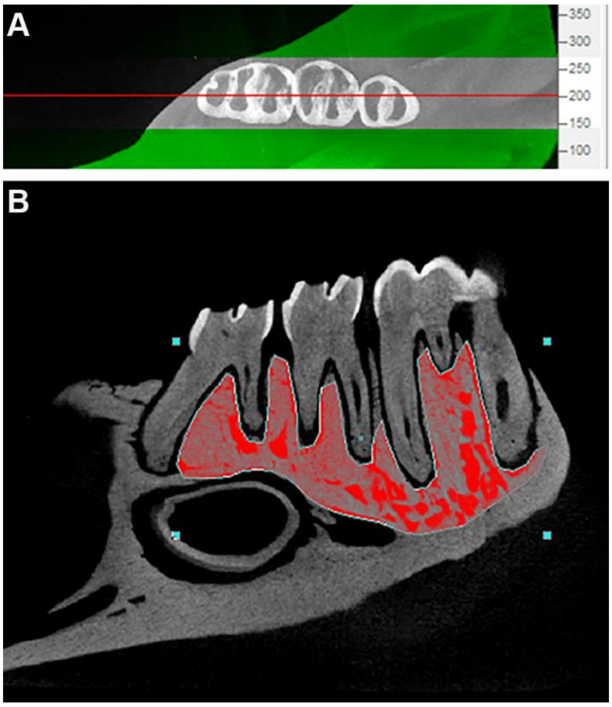

**Supplemental Fig. 2.**

Large hypertrophic bone area from the right side in the BTX21 group. The blue alcian staining evidenced: cartilage with chondrocytes delimited by white dotted line, a transition osteocartilaginous zone delimited by [ ], active remodeling region indicated by *. “Bone” referred to hypertrophic bone area.

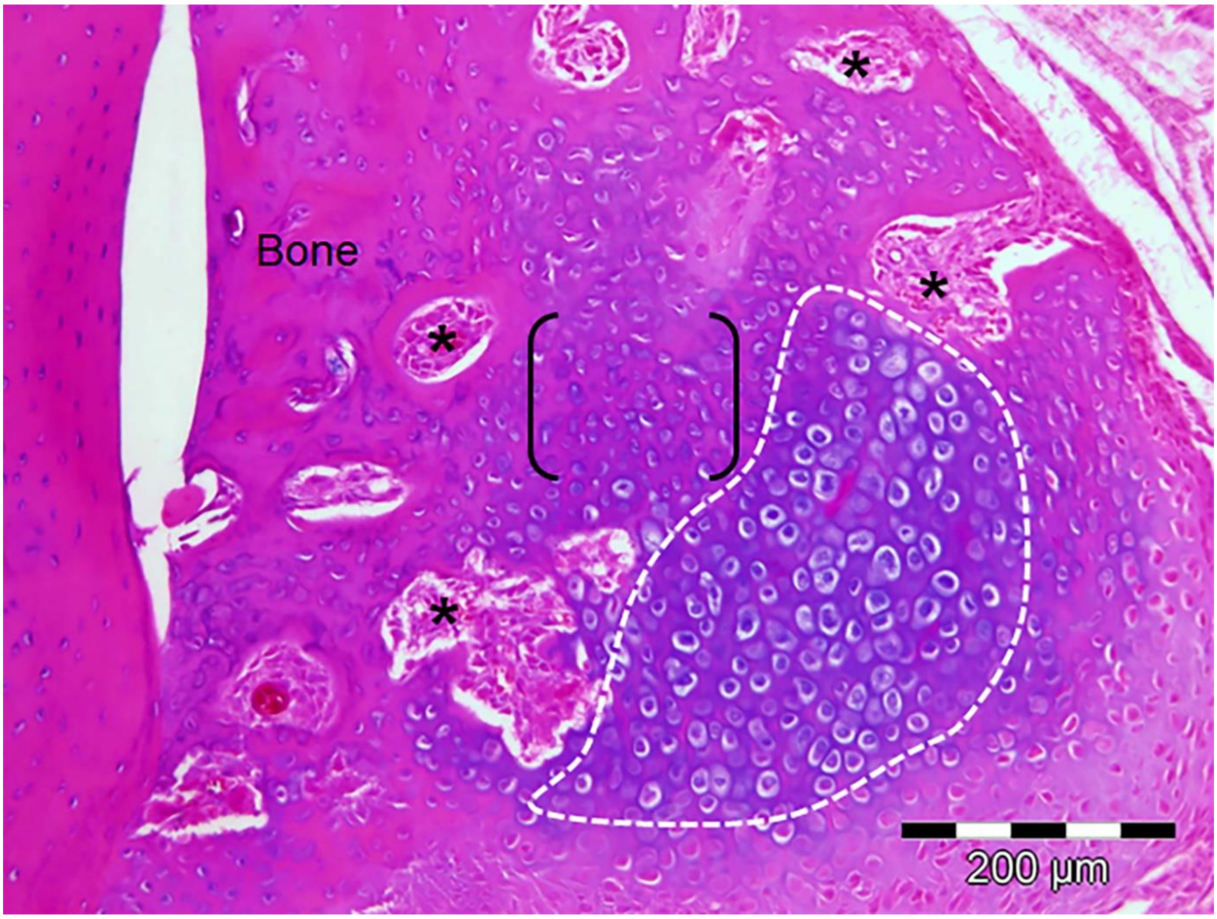
